# A *Mycobacterium tuberculosis* Mbox controls a conserved, small upstream ORF via a translational expression platform and rho-dependent termination of transcription

**DOI:** 10.1101/2025.08.04.667915

**Authors:** Alexandre D’Halluin, Terry Kipkorir, Catherine Hubert, Kristine Arnvig

## Abstract

Magnesium is vital for bacterial survival, and its homeostasis is tightly regulated. Intracellular pathogens like *Mycobacterium tuberculosis* (Mtb) often face host-mediated magnesium limitation, which can be counteracted by upregulating the expression of Mg^2+^ transporters. This upregulation may be via Mg^2+^-sensing regulatory RNA such as the *Bacillus subtilis ykoK* Mbox riboswitch, which acts as a transcriptional “OFF-switch” under high Mg^2+^ conditions. Mtb encodes two Mbox elements with strong similarity to the *ykoK* Mbox.

In the current study, we characterize the Mbox encoded upstream of the Mtb *pe20* operon, which is required for growth in low Mg^2+^ combined with low pH. We show that this switch operates via a translational expression platform and Rho-dependent transcription termination, which is the first such case reported for an Mbox. Moreover, we show that the switch directly controls a small ORF (uORF2) encoded upstream of *pe20*. We have annotated this highly expressed and highly conserved uORF as *rv1805A*, but its role remains unclear. Interestingly, a homologous gene exists outside the Mbox-regulated context, suggesting functional importance beyond magnesium stress.

Overall, this study uncovers a dual mechanism of riboswitch regulation in Mtb, combining translational control with Rho-mediated transcription termination. These findings expand our understanding of RNA-based gene regulation in mycobacteria, with implications for pathogenesis and stress adaptation.

## INTRODUCTION

Magnesium is required for a wide range of cellular functions in all domains of life and the most abundant divalent cation in living cells (Smith *et al*., 1998). In bacteria, these functions include cell wall integrity, biofilm formation, macromolecular metabolism and -function, making magnesium homeostasis essential (Thomas & Rice, 2014; Subramani *et al*., 2016; Yamagami *et al*., 2021; Chatterjee *et al*., 2024). This represents an extra challenge for intracellular pathogens as host immune responses include mechanisms for restricting access to magnesium in certain cellular compartments such as the phagosome (reviewed in Forbes & Gros 2001; Pokorzynski & Groisman 2023). To counteract these defence mechanisms pathogens express an array of transporters to ensure adequate Mg^2+^ uptake. At least four types of magnesium channels and transporters regulate and maintain essential Mg^2+^ levels in prokaryotes: CorA, CorB/C, MgtA/B and MgtE (Franken *et al*., 2022). *Mycobacterium tuberculosis* (Mtb) encodes CorA (Rv1239), MgtE (Rv0362) transporters MgtA/B however, Mtb does not encode homologues of MgtA/B, making the function of MgtC elusive (Alix & Blanc-Potard, 2007).

Riboregulated, i.e. RNA-based, stress-responses are widespread in bacteria, with small RNAs and riboswitches being the most prominent elements. Riboswitches are located in the 5’ leader regions of mRNAs regulating gene expression in *cis*; they are composed of a highly structured ligand binding aptamer domain and an expression platform. The latter exerts gene expression control by either modulating premature termination of RNA Polymerase (RNAP) and/or restricting access of the ribosome to the Ribosome Binding Site (RBS) of the mRNA (Salvail & Breaker, 2023). Nonpermissive control mechanisms may involve the formation of intrinsic terminators, unmasking of Rho-binding sites or occlusion of Shine-Dalgarno (SD) sequence of the downstream Open Reading Frame (ORF). The latter may in addition be associated with Rho-dependent termination of transcription within the ORF (Salvail & Breaker, 2023). Binding of the specific ligand can either allow (“ON-switch”) or inhibit (“OFF-switch”) expression of the downstream gene (Breaker, 2018; Kavita & Breaker, 2023; Schwenk & Arnvig, 2018). The genes regulated by riboswitches are often, but not always, involved in the metabolism or transport of the cognate ligand (Kavita & Breaker, 2023; Roth & Breaker, 2009; Sherlock & Breaker, 2020). Riboswitch ligands range from sugars, amino acids, nucleotides and cofactors to metal ions including Mg^2+^ (Breaker 2022; Barrick *et al*., 2004b; Dann *et al*., 2007; Mccown *et al*., 2017). A magnesium-sensing riboswitch, referred to as Mbox, was first discovered in the *Bacillus subtili*s *ykoK* gene encoding a MgtE-type magnesium transporter (Barrick *et al*., 2004a; Ramesh & Winkler, 2010; Townsend et al., 1995). The Mbox is a transcriptional ‘OFF-switch’; magnesium binding to the aptamer leads to conformational changes of the RNA and the formation of an intrinsic terminator preventing *ykoK* expression. At low Mg^2+^ concentrations, the absence of the terminator is permissive to *ykoK* expression which facilitates increased Mg^2+^ uptake (Ramesh & Winkler, 2010).

Successful infection with Mtb requires its sensing of, and adaptation to, multiple micro-environments including different types of macrophages and their subcellular compartments such as phagosomes (Chandra *et al*., 2022; Samuels *et al*., 2022; Sholeye *et al*., 2022). Mtb has evolved mechanisms to either escape this organelle or to endure the hostile environment within by a range of adaptive responses (Ehrt *et al*., 2018; Ernst, 2012; Huang, 2019). Riboswitches are likely to play a role in this adaptation by directly sensing host environments via specific metabolites. Several Mtb riboswitches have been predicted (Rfam RF00380) and their expression validated by RNA-seq, Term-seq, inline probing and functional assays (Nawrocki *et al*., 2015; Arnvig *et al*., 2011; D’Halluin *et al*., 2023; Kipkorir *et al*., 2024a+b). These include two predicted Mbox aptamers upstream of Rv1535 and Rv1806 (*pe20 locus*), respectively. Although the two Mtb aptamers show a high level of structural similarity to the *Bacillus subtilis ykoK* aptamer, published results suggests that these interact differently with divalent cations including Mg^2+^ (Bahoua *et al*., 2021). The *pe20 locus* (encoding PE20, PPE31, PPE32, PPE33 Rv1810 and MgtC) is associated with magnesium homeostasis and acid stress, and PE20-PPE31 have been shown to be necessary for maintaining growth in a combination of low Mg^2+^ and low pH, conditions that mimic the phagosomal environment (Walters *et al*., 2006; Wang *et al*., 2020). The function of Rv1535 remains unknown.

We recently mapped premature termination of transcription in Mtb at genome-scale and identified hundreds of RNA leaders with an abundance of potential new riboswitches and translated small upstream ORFs (uORFs) (D’Halluin *et al*., 2023). We validated predicted riboswitches and demonstrated that both Mtb Mbox leaders were associated with premature, Rho-dependent termination of transcription upstream of the annotated ORFs (D’Halluin *et al*., 2023).

Here, we show that *pe-ppe* associated Mboxes are widely conserved across *Mycobacteria* and are co-transcribed with *mgtC* in Mtb. The Mtb Mbox upstream of *pe20* is unusual as it combines a translational expression platform with a Rho-dependent transcription terminator. This is to our knowledge the first translational Mbox to be described. Using translational reporter fusion constructs we show that a conserved uORF located between the Mbox and *pe20* is highly expressed. This peptide is highly conserved in the context of Mboxes across the *Mycobacterium* genus. While its function remains opaque, a paralogue of this ORF is expressed from an additional, Mbox-independent *locus* in Mtb, supporting the biological and regulatory importance of this peptide.

## RESULTS

### Conservation and genomic context of *M. tuberculosis* Mboxes

Two Mbox aptamers have been identified within the Mtb H37Rv genome (Rfam RF00380). We used these sequences to predict their structures and compare these to the Mbox consensus structure from Rfam. The results, shown in Figure 1A, indicate a high degree of similarity between the two Mtb aptamers and the *B. subtilis ykoK* element, suggesting these are functional Mg^2+^-sensing elements as reported in the case of *rv1535* by (Bahoua *et al*., 2021).

**Figure 1.**
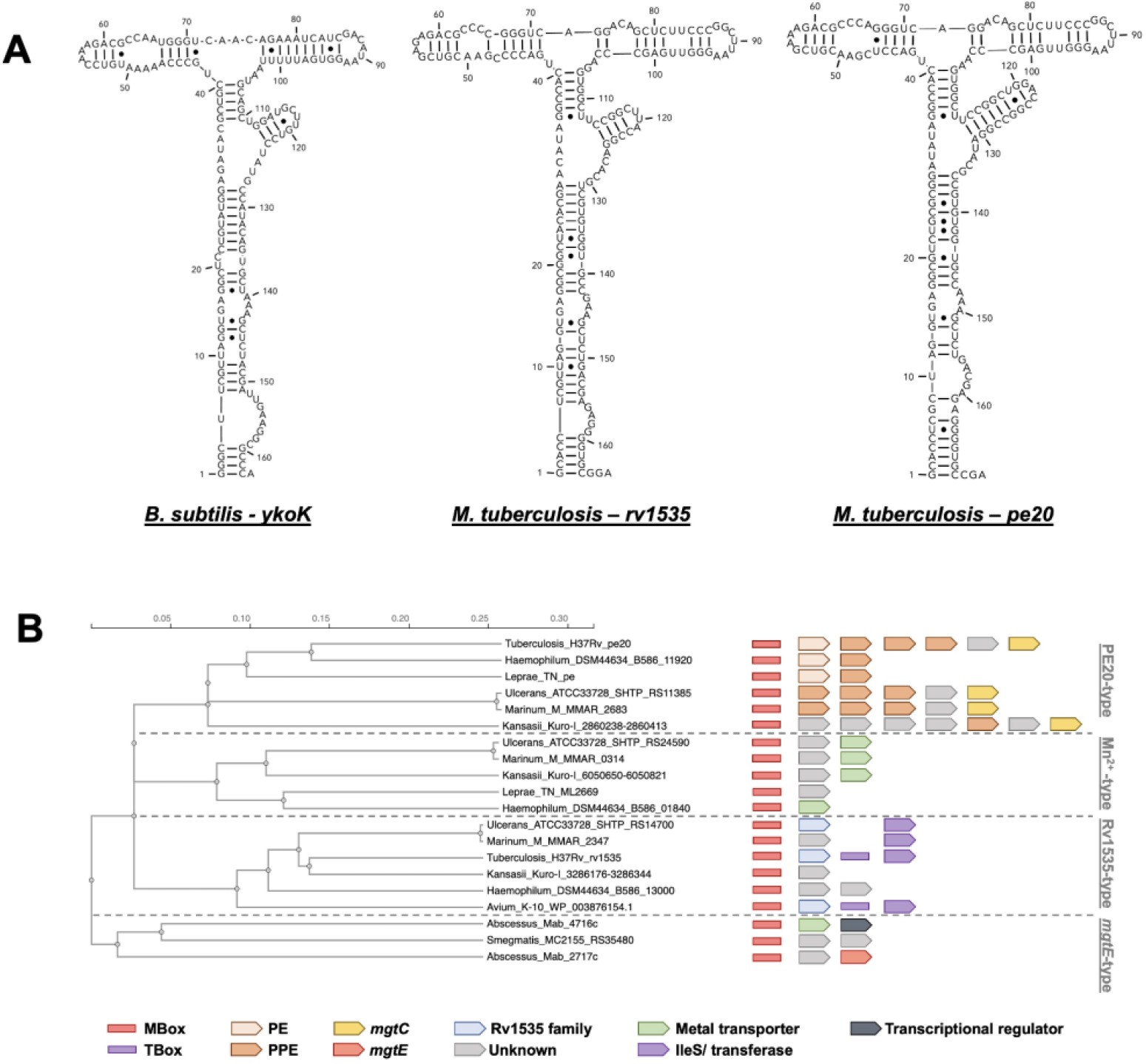
Conservation of Mbox elements. A) Mbox aptamer structures from *Bacillus subtilis ykoK* and the two *M. tuberculosis* aptamers from *rv1535* and *pe20* (*rv1806*). Structures were predicted using RNAstructure Web Serveur (Reuteur & Mathews, 2010) and drawn by extracting the bracket-dot plot to RNAcanvas (Johnson & Simons, 2023). B) Distribution of Mboxes and their associated genes in *Mycobacteria*. These can be split into the four types indicated, based on the aptamer and their downstream sequences. Notably, the two *M. tuberculosis* elements fall into different groups.

Next, we investigated the conservation of the element and its context across the *Mycobacterium* genus. Both elements have been shown to be associated with multiple uORFs, and at least one of these is translated (u2, D’Halluin *et al*., 2023). Based on a phylogenetic analysis of the two Mtb elements and Mboxes from other species, we identified four classes of mycobacterial Mboxes, represented by *pe20*-type, manganese-type (Mn^2+^-type), *rv1535*-type and *mgtE*-type elements, respectively; these classes are further supported by a conserved gene synteny (Figure 1B).

The *mgtE*-type is the only Mbox found in the non-pathogenic *M. smegmatis*, and its genomic neighbourhood shows that the genes immediately downstream encode predicted metal transporters (MT) and/or associated proteins (e.g. MgtE), or proteins of unknown function. The other three branches are seen across fast-and slow-growing pathogenic *Mycobacteria*.

The first branch, the *pe20*-type is almost exclusively found upstream of multiple *pe*-*ppe* genes, which in Mtb, *M. ulcerans, M. marinum* and *M. kansasii* are followed by genes of unknown function (GUF) and MT (as MgtC was originally annotated as a magnesium transporter). The second Mn^2+^-type branch includes a cluster of manganese transporter-type downstream of the riboswitch, a constellation that is not seen in Mtb. The third *rv1535*-type branch is found upstream of GUF like *rv1535* but followed by a cluster of T-box/*ileS* elements or various transferases. These results suggest the regulation of metal transporters by both Mboxes is well-conserved across *Mycobacteria*, while the the *pe-ppe* clusters and mosaic appearance suggest insertion events may have taken place in pathogenic / slow growing species.

### Rho-dependent premature termination of transcription within Mbox leaders

TSS mapping and RNA-seq suggest that the *rv1535* mRNA is monocistronic, while the *pe20* mRNA is polycistronic spanning *pe20* to *mgtC* (Arnvig *et al*., 2011, Cortes *et al*., 2013, D’Halluin *et al*., 2023) (Supplementary figure 1). Importantly, the entire polycistronic *pe20* operon is upregulated during growth in low magnesium controlled by the Mbox (Walters *et al*., 2006).

We recently mapped premature transcription termination (TTS) in Mtb genome-wide and identified two dominant TTS associated with the Mbox leaders (D’Halluin *et al*., 2023). TTS1062 is located ∼210 nucleotides downstream of the *rv1535* TSS and 40 nucleotides downstream of the aptamer. TTS1209 is located ∼185 nucleotides downstream of the *pe20* TSS and 10 nucleotides downstream of the aptamer (Figure 2A).

**Figure 2.**
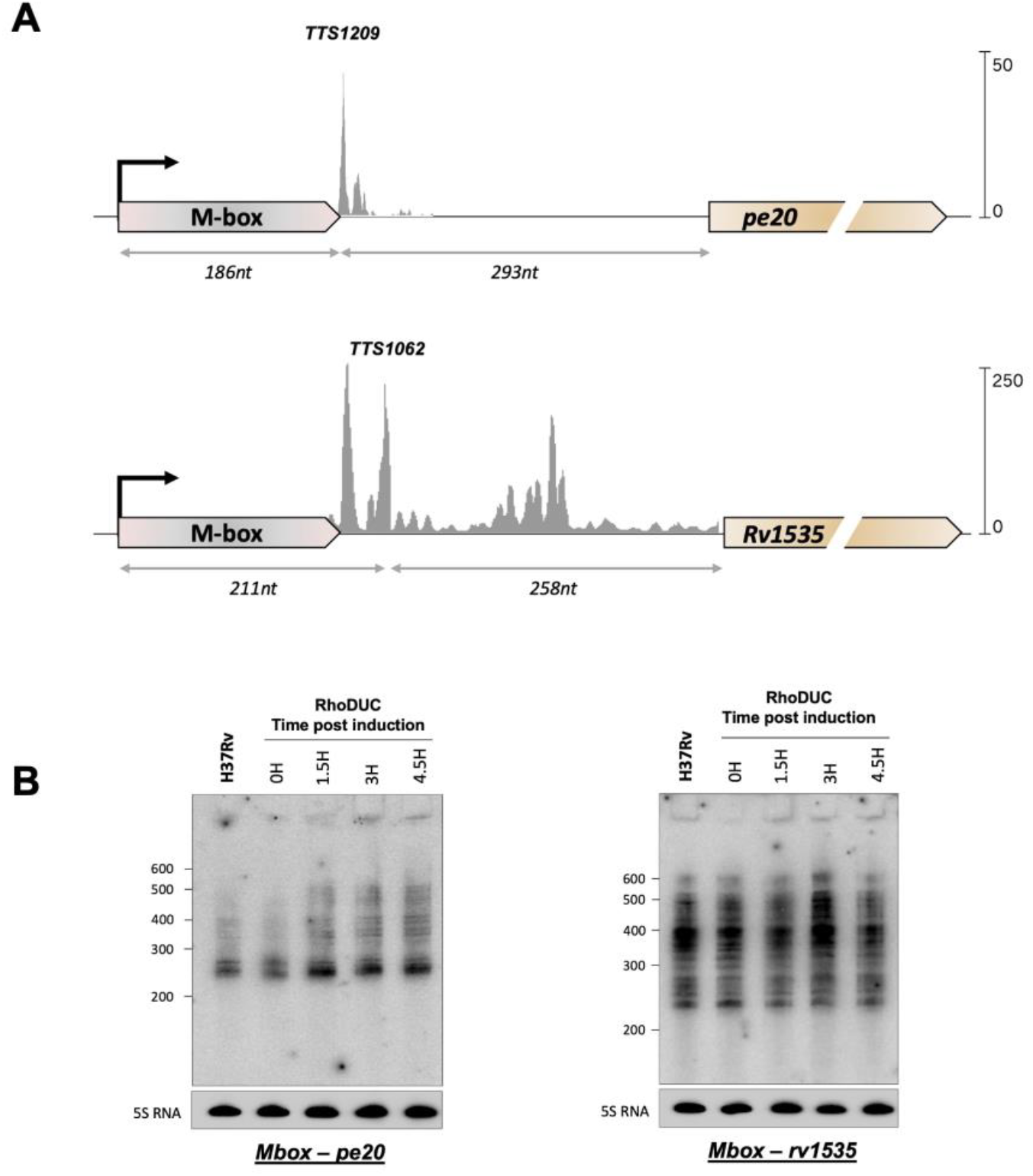
Premature termination of transcription within Mtb Mbox *loci*. A) The two Mbox-associated genes, *rv1535* and *pe20* are shown with their respective leaders. TSS from (Cortes *et al*., 2013), Term-seq data and Transcription termination sites (TTS) from (D’Halluin *et al*., 2023). Distances from TSS to dominant TTS peaks and further to the start codons of have been indicated. B) Northern blot with log-phase total RNA from Mtb H37Rv and from RhoDUC (Botella *et al*., 2017) following depletion of Rho. Total RNA was separated on an 8% acrylamide gel, electroblotted and probed for leader sequences distinct for the two genes, approximately 180 nucleotides downstream of the TSS. The 5S RNA was probed as a loading control.

In both *loci*, multiple smaller peaks are flanking the TTS, suggesting a degree of flexibility in the TTS. Both TTS were located a significant distance (>200 nucleotides) upstream of their annotated ORFs, revealing the premature termination of transcription within the two leader regions, and neither were associated with canonical intrinsic terminator structures.

In our Mtb TTS mapping we predicted and validated Rho-dependent termination using RhoTermPredict (Di Salvo *et al*., 2019) and depletion of Rho using the Rho-DUC strain (Botella *et al*., 2015; D’Halluin *et al*., 2023).

Two Rho-binding (*rut*) sites were predicted in each Mbox leader; one in each aptamer (T5468 and T6425) and one between aptamers and annotated ORFs (T5469 and T6426), while the mapped TTS1209 and TTS1062 are located between these (Table 1; Figure 2A).

**Table 1.** Location of rut sites within the M-box leaders.

| Mbox | TTS position | RTP | rut site end | dist. from TSS | dist. to annotated ORF |
| --- | --- | --- | --- | --- | --- |
| *rv1535* | 1735718 | T5468 | 1735604 | 17 | 372 |
|  |  | T5469 | 1735894 | 307 | 82 |
| *pe20* | 2047779 | T6425 | 2047711 | 38 | 361 |
|  |  | T6426 | 2047965 | 292 | 107 |

The calculated readthrough (RT) scores for the mapped TTS after Anhydrous Tetracyclin (ATc) induced depletion of Rho validated that transcription termination was in fact due to Rho (D’Halluin *et al*., 2023).

To further confirm Rho-dependent, premature termination of transcription, we performed Northern Blotting on RNA from H37Rv and from Rho-depleted cultures probing for both Mboxes. The homology between the Mbox aptamers from *rv1535* and *pe20* made it impossible to design a 5’ probe that could distinguish between the two transcripts. To ensure that the signals were specific for either *pe20* or *rv1535*, we used a probe that was located 180 nucleotides into the transcripts beyond the homologous regions (supplementary figure 2) and as a result, transcripts shorter than this could not be detected. Several strong signals between 200 and 300 nucleotides roughly corresponding to the TTS mapping suggesting multiple points of premature termination of transcription within both leaders (Figure 2B). In H37Rv and in Rho-DUC time 0, we observed limited readthrough beyond 300 nucleotides for *pe20*, while *rv1535* displayed multiple larger signals primarily around 400 nucleotides consistent with the TTS pattern. Depletion of Rho led to an increase in larger transcripts for *pe20* suggesting increased readthrough i.e. reduced termination. In contrast, *rv1535* TTS pattern changed only marginally over the rho depletion time course, suggesting that Rho plays a greater role for *pe20* regulation as compared to *rv1535*.

### The regions downstream of the Mbox aptamers harbour multiple uORFs

Ribosome profiling demonstrates that the regions between the Mbox aptamers and the two annotated open reading frames (ORF)s (*rv1535* and *pe20*, respectively) are bound by ribosomes in agreement with on-going translation upstream of the annotated genes (Sawyer et al., 2021; Smith et al., 2022; D’Halluin et al., 2023). Sequence alignment of *rv1535* and *pe20* leaders with other *pe20*-type leaders indicated several regions of conservation including near-identical Shine-Dalgarno (SD) sequences located at the end of the aptamer (SD1) and a second, highly conserved SD (SD2) further downstream. The ORF downstream of SD1 (upstream ORF1/uORF1) shows poor conservation, while the uORF downstream of SD2 (uORF2) is highly conserved (Supplementary figure 2). Moreover, we have previously shown that uORF2 from both loci is expressed (D’Halluin *et al*., 2023).

To characterize the relationship between the *pe20* Mbox and the two uORFs, we first investigated expression using translational *lacZ*-fusions. All constructs included the 5’ leader from the TSS and were gradually extended downstream to the end of uORF1 (Mbox-*uORF1-lacZ*); the end of uORF2 (Mbox-*uORF2-lacZ*), or the start codon of *pe20* ORF (Mbox-*pe20::lacZ*), respectively. All were fused in-frame to *lacZ*, expressed from a heterologous, constitutive promoter and integrated into the *M. smegmatis* genome in single copy (Figure 3). Next, we performed β-galactosidase (β -gal) assays, which showed that Mbox-*uORF2-lacZ* expression was >10 fold higher than Mbox-*pe20::lacZ* (∼650 Miller Units compared to 60 Miller Units), while Mbox-*uORF1-lacZ* was only slightly higher than the background (Figure 3B). To validate the start codons of the two uORFs, we mutated each to non-start codons (GTG to GTC and ATG to ACG for *uORF1* and *uORF2*, respectively. This reduced expression significantly in both constructs, in support of the suggested translation start sites, although Mbox-*uORF2-lacZ* expression was still higher than Mbox-*pe20::lacZ* expression (Supplementary figure 3).

**Figure 3.**
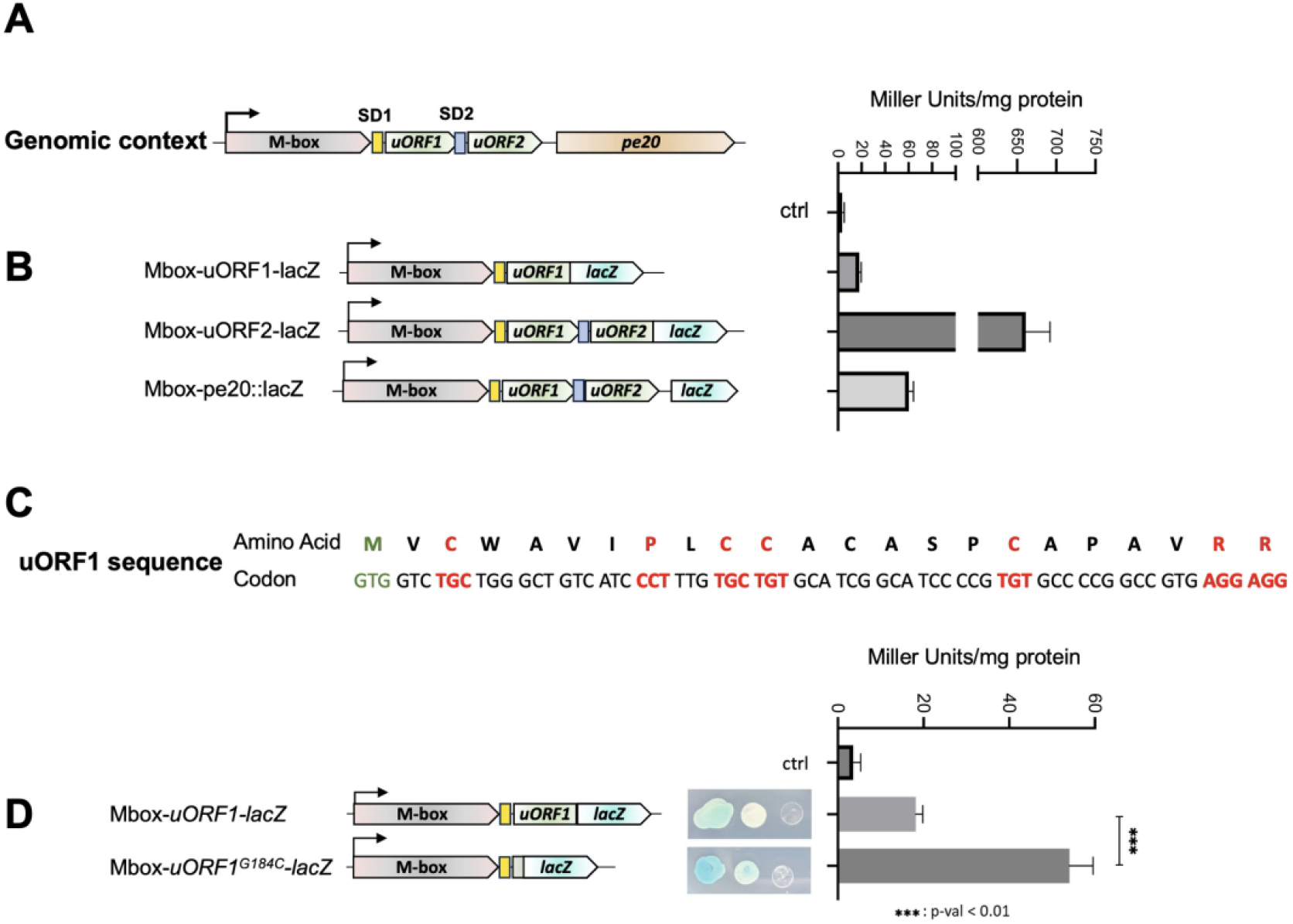
Expression of *pe20* uORFs. To ascertain expression of uORF1 and uORF2 from the *pe20* operon, we made translational *lacZ*-fusions and measured ***β***-galactosidase (***β***gal) activity of the different constructs. Experiments were done in triplicates and differences of expression tested with a t-test (p-val<0.01). A) Genomic context of *pe20* and the uORFs associated (green) including SD1 (yellow box) and SD2 (blue box); B) Schematic showing each reporter constructs (left) and their expression in Miller units (right). C) uORF1 sequence with amino acids and their codons. Start codon is shown in green and rare codons (<5/1000 frequency) are shown in red. D) Reporter constructs with full-length and truncated uORF1 sequence (left) followed by spotting assay of 10 pl of a culture at OD_600nm_ of 0.6, 0.06 and 0.006, indicating expression on Xgal plates and their respective ***β***gal activities are shown on the right.

As the translation initiation region (TIR) for uORF1 and uORF2 (i.e. SD1 and SD2 and their distances to the start codons) were almost identical, and we had not observed any premature TTS in the region, we reasoned that a polypeptide segment of the uORF1 led to lower expression. uORF1 contains several rare (≤5/1000) codons, i.e. TGC, CCT, TGC, TGT, TGT, AGG, AGG (Figure 3C). To explore this possibility, we deleted the majority of uORF1 from the Mbox-*uORF1-lacZ* construct except the first two codons (Mbox-GTG-GTC^uORF1^*-lacZ*) and measured *lacZ* expression.

The results, shown in Figure 3D indicate that expression of this truncated uORF1 was 2.5-fold higher than that of the full-length uORF1, suggesting that the uORF1 sequence did indeed suppress expression.

### The *pe20* Mbox operates via a translational expression platform

In conjunction, the Rho-dependent premature termination of transcription, the highly conserved SDs at the end of the aptamer, located upstream of a well-expressed conserved uORF made us speculate that the *pe20* Mbox operates via a translational expression platform.

A functional translational expression platform requires the potential for the SD to be masked, e.g. by a pyrimidine-rich region (an αSD) that in turn can be sequestered by an ααSD under different conditions.

We identified such a region approximately halfway between SD1 and SD2. This αSD and its flanking regions have the potential to pair with the entire translation initiation region (TIR) of uORF2 (shown in blue in Figure 4) or alternatively, with the aptamer-associated SD1 and its flanks (yellow in Figure 4). To explore this hypothesis further, we measured uORF2 expression after introducing mutations that could interfere with the proposed interactions. One was the abolishing the uORF1 start codon, the rationale being that this would partially unmask SD1 thereby favouring the SD1-αSD interaction, leading to an increase in uORF2-*lacZ* expression

**Figure 4.**
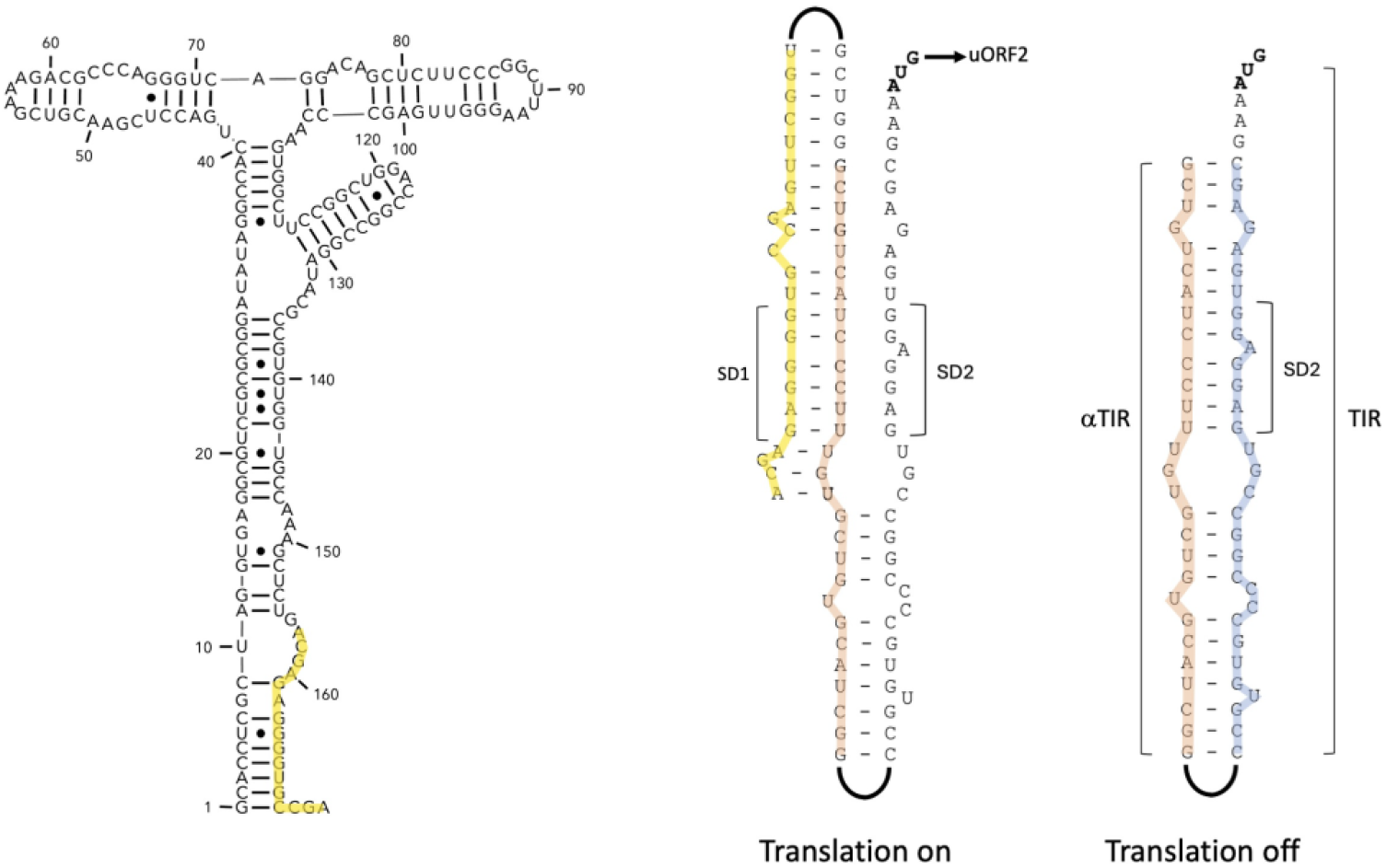
Model for a translational expression platform. The figure shows how the translation initiation region (TIR, blue) can be sequestered by base-pairing with the αTIR (orange), which in turn can base-pair with the ααTIR (yellow), depending on the conformation of the aptamer. Structure of the aptamer is shown on the left with part of the ααTIR shown in yellow.

Similarly, deleting the αSD should also lead to higher expression of uORF2, as SD2 would no longer be sequestered. The results, shown in Figure 5A indicate a moderate (∼1.3-fold), but significant increase in expression, when the start codon of uORF1 was changed (Mbox- *uORF2*^*G184C*^*-lacZ*) and a larger (∼2-fold) increase in expression, when the-αSD was deleted (Mbox-ΔαSD*-uORF2-lacZ*). Combining the two mutations did not result in an additive effect, suggesting they involved the same mechanism (Figure 5A). These results support a model in which SD1 (ααSD), αSD and SD2 interact to control the expression of uORF2-*lacZ*.

**Figure 5.**
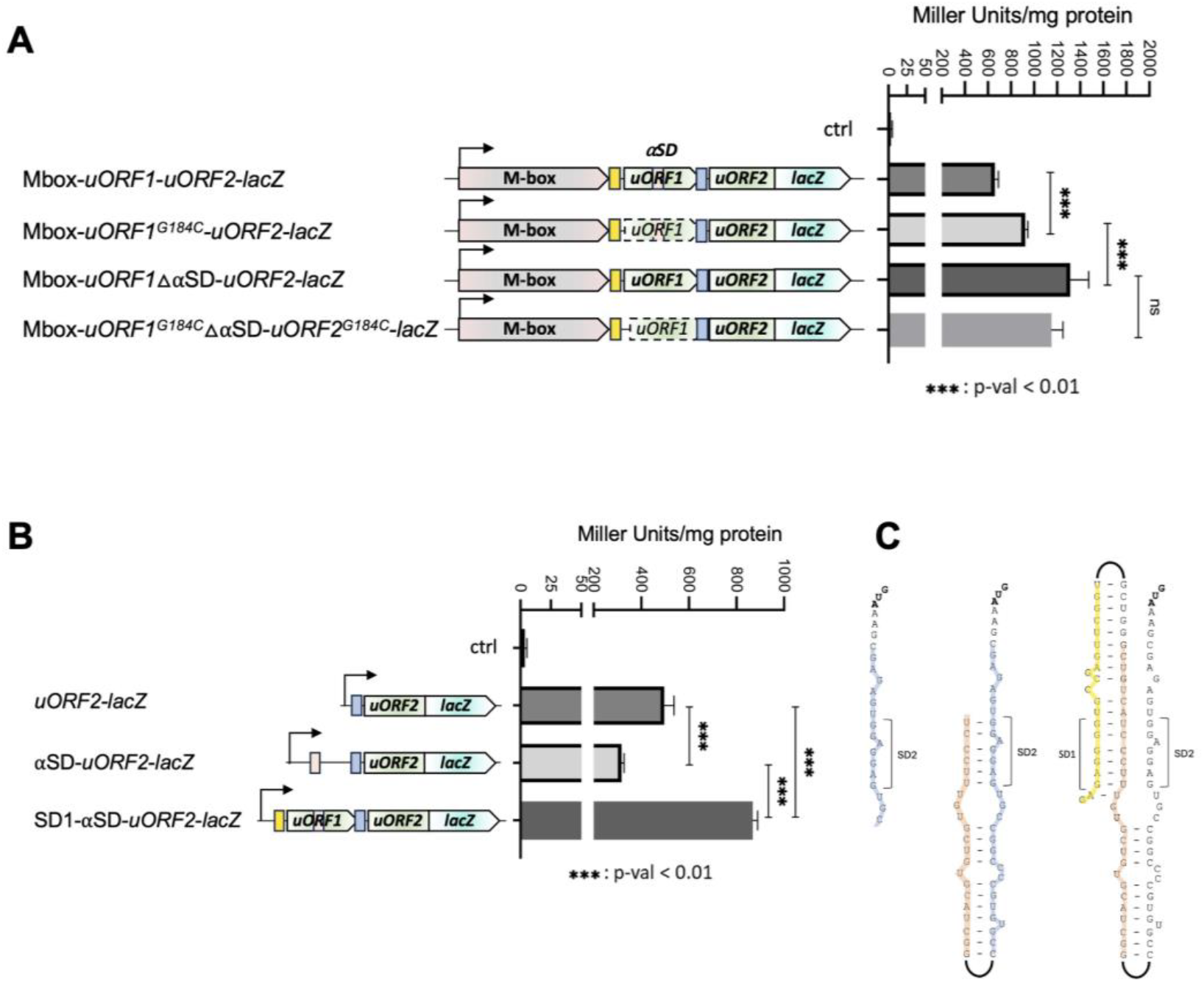
Testing the model for a translational expression platform. A) Reporter constructs assessing the effect of uORF1 changes on uORF2 expression; changing the start codon of uORF1 to a no-start (G184C), deleting the proposed αSD, which is part of uORF1 or a combination of the two. B) Effect of gradual extension of region upstream of uORF2. Expression decreases, when αSD is included and increases again, when SD1 (ααSD) is included. C) Structures indicating how the reporter constructs relate to the model proposed in Figure 4. Experiments were done in triplicates and differences of expression tested with a t-test (p-val<0.01).

To further validate this model, we assessed the contribution of each element by gradually extending the region between uORF2 and the Mbox in uORF2-*lacZ* fusions (Figure 5B). The SD2-uORF2 construct displayed β-gal expression levels of ∼500 Miller Units, and the addition of the αSD motif reduced the β -gal expression by ∼35%. However, a further extension including the ααSD motif led to a substantial increase in uORF2 expression. This is likely due to the unmasking of SD2 and corroborates our model of a translational expression platform controlling uORF2 expression.

Our results support a model in which uORF2 is controlled by a translational Mbox riboswitch combined with Rho-dependent termination of transcription. Based on sequence homology, we propose that the *rv1535* Mbox likewise operates via a translational expression platform. To the best of our knowledge, these are the first examples of an Mbox translational expression platform and Rho-dependent termination of transcription.

### An Mbox-independent homologue of uORF2 encoded in a separate Mtb *locus*

Considering the high conservation between uORF2 in the *rv1535* and *pe20 loci*, we carried out deeper sequence searches and identified a third homologue of the uORF2 region including its SD downstream of the *gca*-*gmhA*-*gmhB*-*hddA* operon. This *locus* is associated with horizontal gene transfer (Becq *et al*., 2007) and the uORF2 homologue annotated as Rv0115A (Figure 6A).

**Figure 6.**
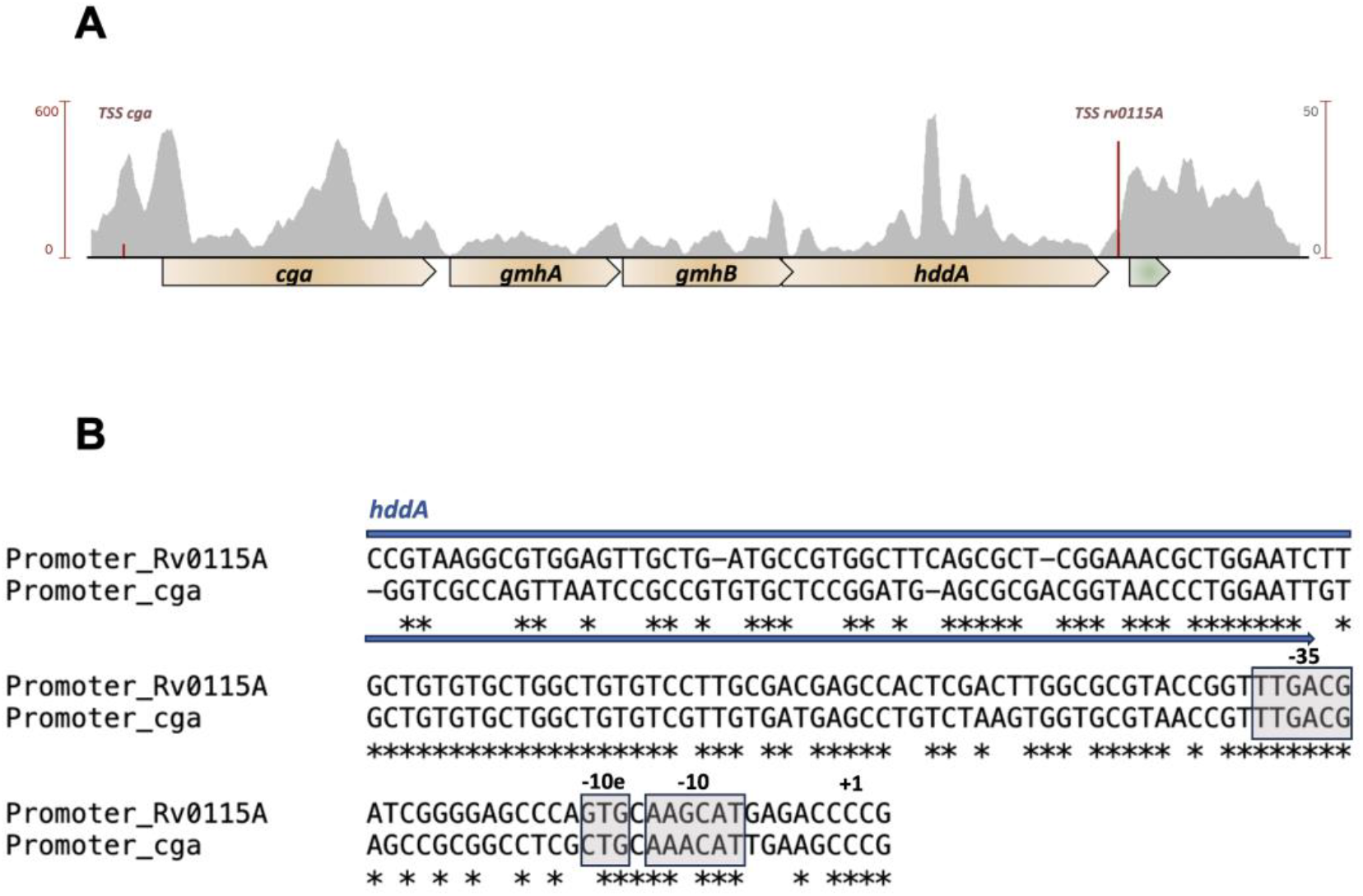
The RvO115A locus. BLAST identified RvO115A to be a homologue of Rv1805A. A) RvO115A (green) is encoded downstream of the *gca* operon (golden) but transcribed from its own promoter. B) Alignment of the promoter regions of *gca* and *rvO115A* show high degree of similarity, suggesting a duplication event. The blue arrow indicates *hddA* coding sequence upstream of *rvO115A*. The promoter elements, −35, extended −10 (−10e) and −10 are highlighted in grey.

We identified two TSS and associated promoter motifs within this *locus*. The first drives the transcription of the *gca-hddA* operon, which terminates downstream of *hddA* (D’Halluin *et al*., 2023). The second drives the transcription of *rv0115A*, and potentially also a second ORF, *rv0115B*. The *gca* and *rv0115A* promoters have similar unusual motifs in the form of an AANCAT −10 hexamer, an extended −10 motif (TGN), a perfect −35 hexamer and in the case of *cga*, a Cytidine TSS (Figure 6B). A further alignment of the promoter regions from −120 to a few basepairs downstream of the mapped TSS, had a remarkable similarity more than 100 basepairs upstream of the TSS suggestive of a gene duplication event (Figure 6B).

Alignment of uORF2 homologues including *rv0115A* across mycobacterial species reveals a well-conserved N-terminal domain, including a universally conserved Proline residue (Figure 7). This peptide is specific for Mycobacteria, which indicates that uORF2 peptides and their homologues have functions uniquely associated with this genus. Based on this finding, we suggest renaming uORF2 from the *pe20* operon *rv1805A*.

**Figure 7.**
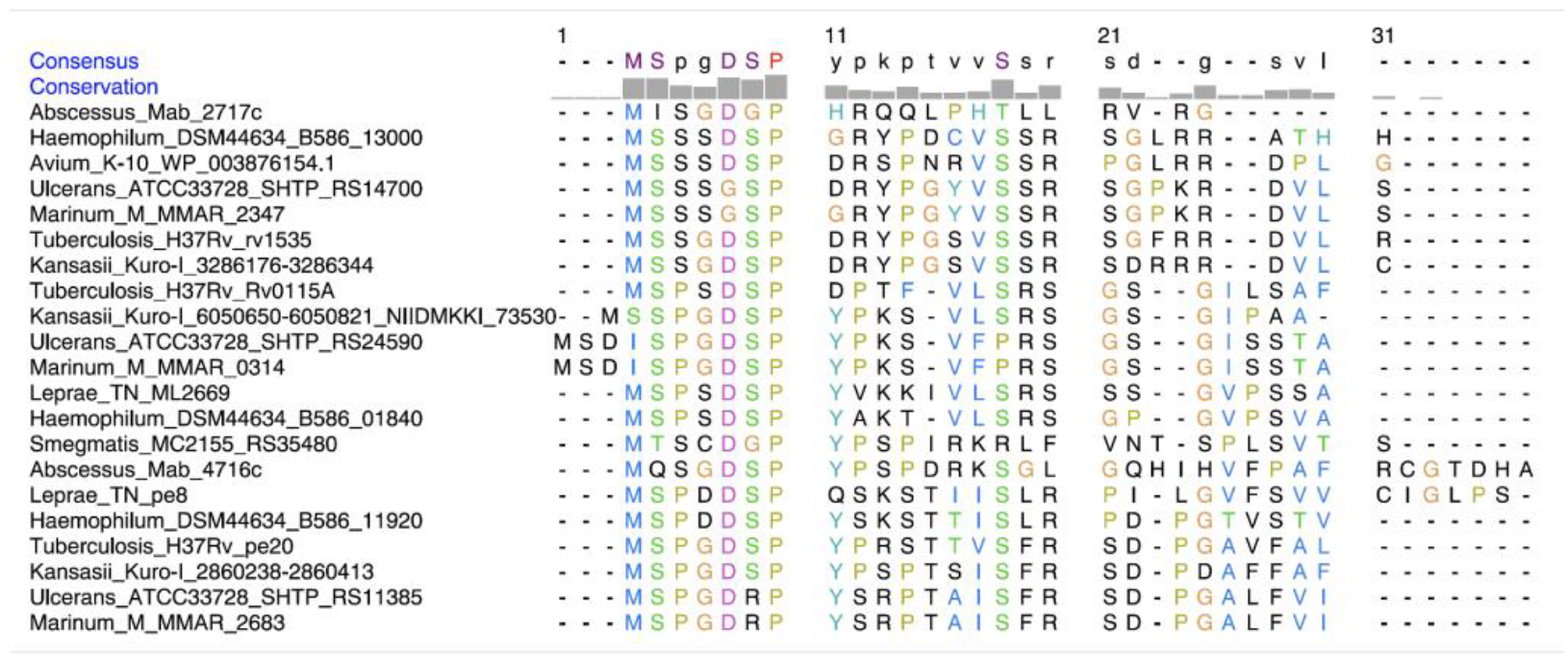
Conservation of uORF2 within Mycobacteria. Alignment of Mbox associated uORF2 extracted from Figure 1B and Mtb Rv0115A peptides showed high conservation of several residues mainly at the N-terminal sequence, including 100% conservation of a proline at position 7 in most peptides. Consensus sequence and amino acid conservation were assessed using Chimera (Meng *et al*., 2006).

### No evident role for Rv1805A in biofilm formation during magnesium stress

Realising the ubiquitous presence of Rv1805A homologues, we sought to find a role for this peptide. PE20 and PPE31 are necessary for Mtb growth in conditions of low Mg^2+^ combined with low pH (Wang *et al*., 2020). To probe a potential role of uORF2 in this process, we exploited the fact that magnesium is required for *Mycobacterium* biofilm formation (Chatterjee *et al*., 2024) and leveraged the trick that *Mycobacterium smegmatis*, a closely related species, has no homolog of *pe20* locus.

In agreement with literature, the growth and biofilm formation of *M. smegmatis* were compromised in low Mg^2+^, and that this phenotype was exacerbated at acidic pH values (Figure 8). We tested whether the expression of *pe20-ppe31* or *rv1805A-pe20-ppe31* might rescue this phenotype by transforming *M. smegmatis* with plasmids expressing the cognate genes. The results in figure 8 indicate no visible difference between strains expressing *pe20-ppe31* with or without *rv1805A* or *rv0115A*; further investigations are required to identify a role of this peptide and its homologues in mycobacterial biology.

**Figure 8.**
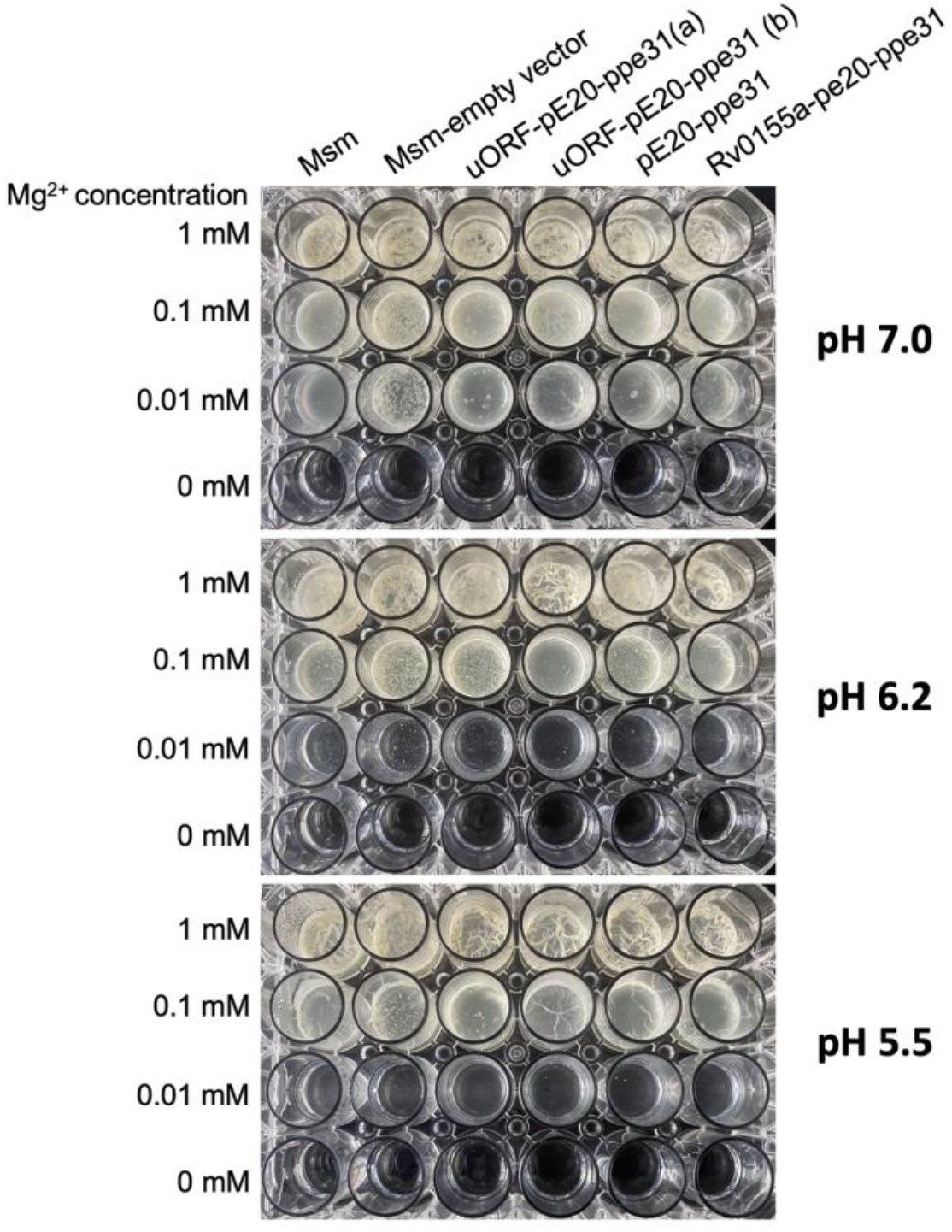
Biofilm formation in *M. smegmatis* during Mg2+-depletion and acid stress. Cultures of *Mycobacterium smegmatis* were grown to mid-log phase, washed in Mg2+-free medium, resuspended in 1 mL of indicated medium at OD 0.01. Plates were sealed in plastic bags and left for static incubation at 37°C_ _ for a week. Plates shown are representative of three independent experiments.

## DISCUSSION

In the current study we have discovered a novel complex magnesium-controlled riboregulatory system which controls *pe20* gene expression in Mtb. Our results show that premature termination occurring in the 5’ leader of *pe20* (and *rv1535*) relies on Rho-dependent termination of transcription (D’Halluin *et al*., 2023). Moreover, the *pe20* Mbox contributes a translational expression platform, where the translation initiation region including the SD of the first gene in the operon can be sequestered by an anti Shine-Dalgarno motif (α SD). This is also, to the best of our knowledge, the first example of a translationally controlled Mbox. This type of control is consistent with the scarcity of intrinsic terminators in Mtb, and it echoes the finding that a mycobacterial T-box is the only known T-box with a translational expression platform (Sherwood *et al*., 2018). Finally, we identified a highly conserved uORF (*rv1805A*), which is the primary regulated ORF within the *pe20* operon and, based on homology, likely also in the *rv1535* operon.

The *pe20* operon is suppressed in the presence of high Mg^2+^ like the Mbox controlled *ykoK* gene in *B. subtilis* (Walters *et al*., 2006; Ramesh & Winkler, 2010). *pe20* and *ppe31* are critical for magnesium uptake in low-pH/low-magnesium conditions suggesting that the gene products form (part of) a magnesium transporter (Feng e*t al*., 2021; Wang *et al*., 2020). We propose that the Mbox–*rv1805A* module acts as the key regulatory gate, enabling expression of the magnesium-responsive PE/PPE transporter complex only under specific environmental conditions, such as low Mg^2+^ and acidic pH.

The structure of the *pe20* operon, including the presence of *mgtC* raises questions about its ancestry. Given what is known about *pe-ppe* gene expansion (Fishbein *et al*., 2015) and what we have observed in other riboswitch-controlled *pe/ppe loci (i*.*e. the Cbl-ppe2-cobQ locus*, and the PE-containing uORF recently identified downstream of the Mtb glycine riboswitch (D’Halluin *et al*., 2023; Kipkorir *et al*., 2024), it is tempting to speculate that an early *pe*(−*ppe*) element invaded the current *pe20 locus* and subsequently expanded whereby Rv1805A became the first gene in this operon.

A recent study suggests that the *rv1535* Mbox, and by extension likely also the *pe20* Mbox associates with other divalent cations in addition to Mg^2+^ (Bahoua *et al*., 2021). Regardless of the identity of the cognate ligand, our results suggest an ability to alternate between two structures: a non-permissive (ligand-bound) structure that sequesters SD1, allowing αSD/ αTIR to pair with SD2/TIR thereby preventing translation of uORF2/Rv1805A. This could in turn lead Rho-dependent termination of transcription, which will affect the entire operon (Hao *et al*., 2021; Molodtsov *et al*., 2023). We note, however, that according to Term-seq results, the primary TTS is located upstream of *rv1805A*, suggesting that Rho-dependent termination does not depend on translation of this uORF. An alternative explanation of our results could therefore be that the pyrimidine-rich region that we have annotated as α SD, might act as a Rho-binding (*rut*) site that would be masked by translation of uORF1. Deleting this region increased expression 2.5-fold, likely due to reduced termination of transcription or by unmasking of SD2. The marginal increase in the expression of uORF2 in the context of an untranslated uORF1 (Mbox-*rv1805A*^*G184C*^*-lacZ*, Figure 5) and the conservation of the ααSD-αSD interaction, suggests a functional interaction. The two models are not mutually exclusive, and further experiments will eludidate the structural and mechanistic basis underlying the riboregulation..

What is the function of Rv1805A? Given its high conservation and position upstream of *pe20*, we hypothesize that Rv1805A acts as a regulatory peptide modulating the activity or assembly of the PE20–PPE31 complex. Alternatively, it may serve as a structural component of a magnesium-responsive transporter. Conservation between Rv1805A, the Rv1535 uORF2 (Rv1535A) and Rv0115A suggests important roles for these peptides and future work will focus on identifying interaction partners of Rv1805A and assessing its impact on magnesium uptake and stress responses.

In conclusion, our findings reveal a previously unrecognized mode of riboswitch control in Mtb, where a translational Mbox integrates with Rho-dependent termination to regulate a conserved uORF. This multilayered modus operandi underscores the sophistication of RNA-based regulation in Mtb stress adaptation.

## MATERIAL AND METHODS

### Strains and cultures

Strains used in this study are listed in Supplementary Table 1. *M. tuberculosis* H37Rv and *M. smegmatis* MC^2^ 155 were cultured on solid media Middlebrook agar 7H11 supplemented with 10% OADC (Sigma), 0.5% Glycerol and 50 µg/ml hygromycin if appropriate. Liquid cultures were done in Middlebrook 7H9 supplemented with 10% ADC (Sigma), 0.5% Glycerol, 0.05% Tween 80 and 50 µg/ml hygromycin where appropriate. Cultures were harvested at an OD_600nm_∼0.6 for mid-log phase.

Mtb RhoDUC strain, a gift obtained from Pr. Dirk Schnappinger, was grown as previously described with 50 µg/ml hygromycin, 20 µg/ml Kanamycin and 50 µg/ml zeocin (Botella *et al*., 2017; D’Halluin *et al*., 2023). When the cultures reached an OD_600nm_∼0.6, depletion of Rho was induced using 500 ng/ml of anhydrotetracycline. Cells were harvested after 0, 1.5, 3 and 4.5 hours.

*Escherichia coli* DH5α was used for cloning the *lacZ* fusion reporters and were cultured on solid LB 1.5% agar supplemented with 50 µM of 5-bromo-4-chloro-3-indolyl-β;-D-galactopyranoside (X-gal) or in liquid LB supplemented with 250 µg/ml Hygromycin.

### Plasmids constructions and primers

Plasmids and primers used in this study are listed in Supplementary Table 1 and 2. pIRATE plasmids, describe in D’Halluin *et al*., 2023, were used for *lacZ* translational fusion reporters and for Beta-galactosidase assay. Reporters were constructed using Gibson assemblies with oligos (Sigma) or geneBlocks (IDT) listed in Table 3 between HindIII and NcoI sites. Start codons point mutations and deletion of the αSD sequence were generated using the Q5 Site-Directed Mutagenesis Kit (New England Biolabs). Plasmids were cloned in *E. coli* DH5α, extracted and sequenced by Sanger sequencing. Plasmids were transformed into *M. smegmatis* by electroporation and selected on Middlebrook 7H11 agar plates containing 50 µg/ml Hygromycin.

### RNA extraction and Northern Blotting

*M. tuberculosis* H37Rv were stopped using 37.5% of cold ice and centrifuge 10min at 5000 rpm 4°C. Total RNA were extracted as previously described using the FastRNA Pro Blue kit (MP Biomedicals) according to the manufacturer’s protocol (Arnvig *et al*., 2011; D’Halluin *et al*., 2023). RNA concentration and purity was assessed using the Nanodrop 2000 (ThermoFisher), residual genomic DNA removed using Turbo DNase (ThermoFisher) and RNA integrity assessed with 2100 Bioanalyzer (Agilent). 10 µg of total RNA were separated on a denaturing 8% acrylamide:bis-acrylamide (19:1) gel and transfer to a nylon membrane. An RNA probe was synthetized using the mirVana miRNA probe synthesis kit (Ambion) to reveal the *pe20* and *rv1535* Mbox transcripts and labelled with 3µM final concentration of ^32^P α-UTP (3000Ci/mmol; Hartmann AnalyticGmbH). Northern blots were revealed using radiosensitive screens and visualized on a Typhoon FLA 9500 phosphoimager (GEHealthcare).

### Beta-galactosidase activity

*M. smegmatis* carrying the *lacZ* reporter fusions were cultured at OD_600nm_ ∼0.6 and centrifuge 10min 5000 rpm. Pellets were washed four times in Z-buffer composed of 60mM Na_2_HPO_4_, 40mM NaH_2_PO_4_, 10mM KCl, 1mM MgSO_4_ and lysed using beads with the FastPrep bio-pulveriser (MP Biomedicals). The supernatant was kept after centrifugation and the protein level assessed using a Bradford yield with the BCA kit (ThermoFisher) following the manufacturer’s recommendations. Beta-galactosidase were done using the Beta-galactosidase assay kit (ThermoFisher) following manufacturer’s protocol. Proteins were pre-incubated for 5min at 28°C before addition of ONPG.

### Biofilm formation

*M. smegmatis* expressing *pe20-ppe31, rv1805A-pe20-ppe31* or *rv0115A+pe20-ppe31* was grown to mid-log phase, washed in Mg^2+^-free medium, resuspended in 1 mL of the indicated medium at OD 0.01 and seeded in 24-well plates. Plates were sealed in plastic bags and left for static incubation at 37°C for a week. Biofilm formation was monitored every day for a week.

### Folding, sequence conservation and distribution across *Mycobacteria*

Representative genomes of several *Mycobacteria* were selected for sequences conservation: *Mycobacterium tuberculosis* H37Rv (NB_000962), *Mycobacterium leprae* TN (AL450380), *Mycobacterium avium* K10 (NZ_CP106873), *Mycobacterium kansasii* Kuro I (AP023343), *Mycobacterium ulcerans* ATCC33728 (NZ_AP017624), *Mycobacterium marinum* M (CP000854), *Mycobacterium abscessus* ATCC19977 (NC_010397), *Mycobacterium* haemophilum DSM 44634 (CP011883) and *Mycobacterium smegmatis* MC^2^ 155 (NZ_CP009494).

The aptamer sequences of the Mboxes were extracted from RFam database (Rfam RF00380) (Nawrocki *et al*., 2015) and extended to the next annotated ORF. DNA and peptidic sequences were aligned using ClustalW (Thompson *et al*., 1994), and alignment strengthen using T-coffee (Notredame *et al*., 2000). The conservation of uORF2 within *Mycobacteria* was determined using Blast (Altschul *et al*., 1990) and amino acid sequences aligned using ClustalW (Thompson *et al*., 1994) and Chimera (Meng *et al*., 2006). The phylogenetic tree was generated by Clustal Omega using the sequences from the aptamer sequence to the start codon of the next in frame annotated ORF (Sievers & Higgins, 2021). Aptamer secondary structures were predicted using the RNAstructure Web Serveur for RNA Secondary Structure Prediction (Reuter & Mathews, 2010). The resulting Connectivity Table (CT) file was then uploaded to RNAcanvas (Johnson & Simon, 2023) for visualization and structural editing

## AUTHOR CONTRIBUTIONS

AD, TK, KBA designed the study. AD, TK, CH, KBA performed experiments. AD, TK, KBA performed data analysis and wrote the manuscript.

## ACKNOWLEDGEMENTS

We are grateful for Finn Werner’s comments on the mannuscript

