## Supplementary material for "A *Mycobacterium tuberculosis* Mbox controls a conserved, small upstream ORF via a translational expression platform and rho-dependent termination of transcription"

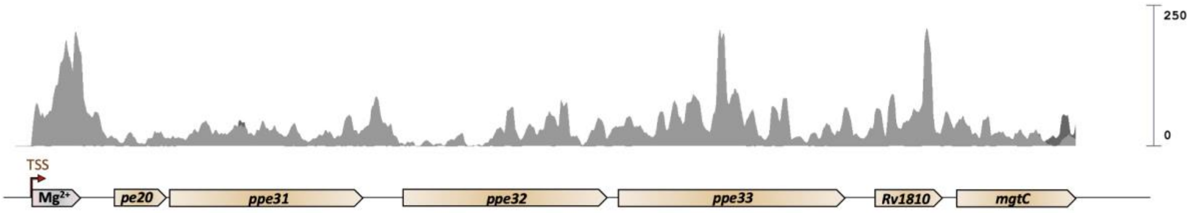


**Supplementary figure 1: The *pe20-mgtC* operon.** RNA-seq traces from Arnvig *et al.*, 2011 were used to map the *pe20* Mbox transcript (grey box, $Mg^{2+}$) which extend to *pe20-ppe31-ppe32-ppe33-rv1810-mgtC* (golden).

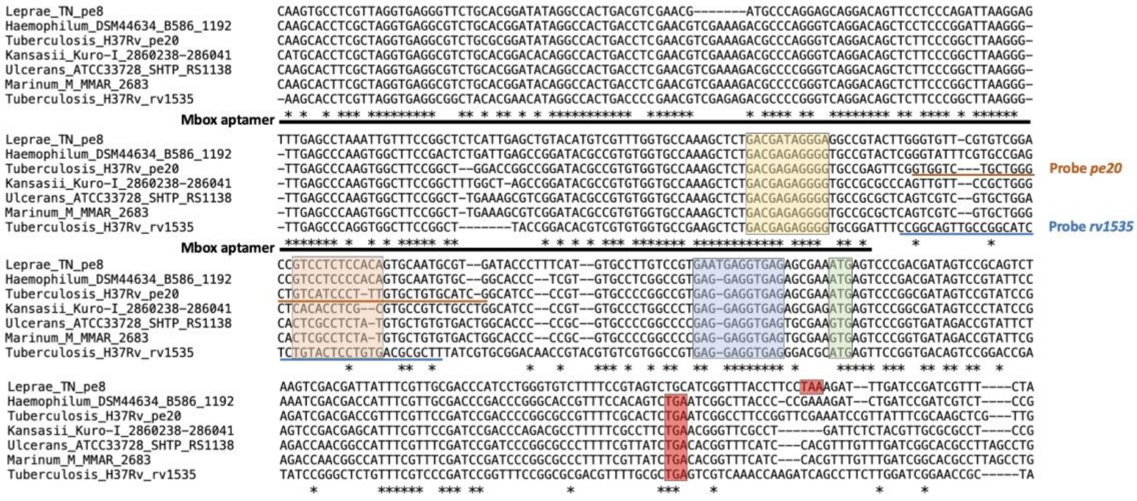


**Supplementary figure 2: Conserved sequences from *Mycobacterial* Mbox aptamer and expression platform.** The sequences from the TSS to the start codon of the annotated ORF of *pe20*-type Mbox from figure 1B and *rv1535* were aligned, highlighting two conserved potential SD (yellow and blue boxes) and a pyrimidine-rich region (orange). A conserved ATG (green) was found downstream of the second SD, alongside a quite conserved stop codon (red). The Mbox aptamer is annotated as a black bar. The probes used for northern blotting are shown for *pe20* (orange bar) and *rv1535* (blue bar).

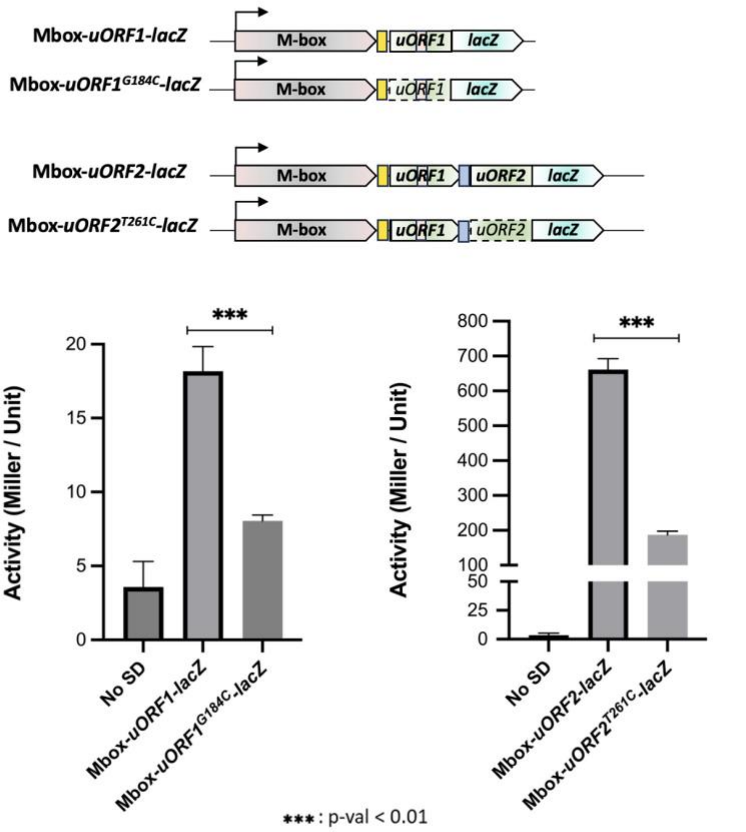


**Supplementary figure 3: Start codon validation of pe20 associated uORFs.** The start codon of uORF1 and uORF2 were respectively mutated from GTG to GTC (G184C) and from ATG to ACG (T261C) in our reporter system (Top). ***β***galactosidase production was measured from triplicates (Bottom) of mutated and non-mutated reporters, and the significance of activity difference assessed with a t-test.

### Supplementary Table 1: Strains and plasmids used in this study

| Strains and plasmids | Description | Reference |
|---|---|---|
| **Strains** | | |
| *M. tuberculosis* | | |
| H37Rv | Virulent model organism of *M. tuberculosis* | Cole et al., 1998 |
| RhoDUC | Δ*rho*::*hygR*, att-L5::rev*tetR*-PtetOFF-*rho*DAS-*zeoR*, Tweety::*tetR*-PtetOFF-*sspB*-*kanR*. Rho-inducible deletion strain with anhydrous tetracyclin. | Botella et al., 2017 |
| *M. smegmatis* | | |
| MC2 155 | Laboratory strain of *M. smegmatis* | Snapper et al., 1990 |
| *E. coli* | | |
| DH5α | Cloning strain for plasmids construction | NEB (C2987) |
| **Plasmids** | | |
| pCR-Blunt II-TOPO | Cloning vector | ThermoFisher (K280002) |
| pIRaTE2020 | Expression vector carrying the *lacZ* ORF under the *rrnB* promoter | D'Halluin et al., 2023 |
| pIRaTE2020_Mbox uORF1-*lacZ* | pIRaTE2020 containing the *pe20* Mbox and the full uORF1 fused to *lacZ* | This work |
| pIRaTE2020_Mbox GTG-GTC$^{uORF1}$-*lacZ* | pIRaTE2020 containing the *pe20* Mbox and the fisrt two codon of uORF1 fused to *lacZ* | This work |
| pIRaTE2020_Mbox uORF1$^{G184C}$-*lacZ* | pIRaTE2020 containing the *pe20* Mbox and the start codon mutated version of the full uORF1 fused to *lacZ* | This work |
| pIRaTE2020_Mbox uORF2-*lacZ* | pIRaTE2020 containing the *pe20* Mbox and the full uORF2 fused to *lacZ* | This work |
| pIRaTE2020_Mbox uORF2$^{T261C}$-*lacZ* | pIRaTE2020 containing the *pe20* Mbox and the start codon mutated version of the fulll uORF2 fused to *lacZ* | This work |
| pIRaTE2020_Mbox uORF2$^{G184C}$-*lacZ* | pIRaTE2020 containing the *pe20* Mbox, the full uORF2 fused to *lacZ* and the start codon of uORF1 mutated | This work |
| pIRaTE2020_Mbox △αSD-uORF2-*lacZ* | pIRaTE2020 containing the *pe20* Mbox, the full uORF2 fused to *lacZ* and the deletion of the αSD sequence | This work |
| pIRaTE2020_Mbox △αSD-uORF2$^{G184C}$-*lacZ* | pIRaTE2020 containing the *pe20* Mbox, the full uORF2 fused to *lacZ*, the deletion of the αSD sequence and the start codon of uORF1 mutated | This work |
| pIRaTE2020 uORF2-*lacZ* | pIRaTE2020 containing the SD$^{uORF2}$ and the full uORF2 fused to *lacZ* | This work |
| pIRaTE2020 △αSD-uORF2-*lacZ* | pIRaTE2020 containing the αSD sequence, the SD$^{uORF2}$ and the full uORF2 fused to *lacZ* | This work |
| pIRaTE2020 SD$^{uORF1}$-△αSD-uORF2-*lacZ* | pIRaTE2020 containing the SD$^{uORF1}$, the αSD sequence, the SD$^{uORF2}$ and the full uORF2 fused to *lacZ* | This work |

| | | |
|---|---|---|
| pIRaTE2020_Mbox *pe20*::*lacZ* | pIRaTE2020 containing the *pe20* Mbox to the start codon of *pe20*, replaced by *lacZ* | This work |

### **Supplementary Table 2 :** Primers and Gblocks used in this study

**Primers**

| Name | Sequence | Use |
|---|---|---|
| Mbox *pe20* | GTGGTCTGCTGGGCTGTCATCCCTTTGTGCTGTGCATC GGCCTGTCTC | Northern Blot probe |
| Mbox *rv1535* | CCGGCAGTTGCCGGCATCTCTGTACTCCTGTGACGCGC TTGCCTGTCTC | Northern Blot probe |
| pIR_F | TTGACTCCATTGCCGGAT | PCR to amplify the mutated inserts from PCR2.0 |
| pIR_R | GACGTTGTAAAACGACGGGA | PCR to amplify the mutated inserts from PCR2.0 |
| Fusion_NoRBS_F | GCCGGATTTGTATTAGACTAAGCTTGAGTAGGAGATT TTCACCTCCTTTCCTTCCTACCATGGATGATCCCGTCGT TTTACA | Oligo Annealing and Gibson Assembly |
| Fusion_NoRBS_R | TGTAAAACGACGGGATCATCCATGGTAGGAAGGAAA GGAGGTGAAAATCTCCTACTCAAGCTTAGTCTAATACA AATCCGGC | Oligo Annealing and Gibson Assembly |
| uORF1-G184C_F | GAGTTCGGTCGTCTGCTG | Q5 mutagenesis |
| uORF1-G184C_R | GGCACCCCTCTCGTCAGA | Q5 mutagenesis |
| uORF2-T261C_F | AGCGAAACGAGTCCCGGCGA | Q5 mutagenesis |
| uORF2-T261C_R | CTCACCTCCTCACGGCCG | Q5 mutagenesis |
| △αSD_F | GTGCTGTGCATCGGCATCC | Q5 mutagenesis |
| △αSD_R | TGACAGCCCAGCAGACCACC | Q5 mutagenesis |

**Gblocks**

| Name | Sequence |
|---|---|
| SD2-uORF2 | GCCGGATTTGTATTAGACTAAGCTTCCGTGAGGAGGTGAGAGCGAAATGAGTCCCGGCGATA GTCCGTATCCGAGATCGACGACCGTTTCGTTCCGATCCGACCCCGGCGCCGTTTTCGCACCCAT TGGATGATCCCGTCGTTTACA |
| antiSD-SD2-uORF2 | GCCGGATTTGTATTAGACTAAGCTTTCCCTTTGTGCTGTGCATCGGCATCCCCGTGTGCCCCGG CCGTGAGGAGGTGAGAGCGAAATGAGTCCCGGCGATAGTCCGTATCCGAGATCGACGACCG TTTCGTTCCGATCCGACCCCGGCGCCGTTTTCGCACCCATTGGATGATCCCGTCGTTTACA |
| SD1-antiSD-SD2-uORF2 | GCCGGATTTGTATTAGACTAAGCTTGAGAGGGGTGCCGAGTTCGGTGGTCTGCTGGGCTGTC ATCCCTTTGTGCTGTGCATCGGCATCCCCGTGTGCCCCGGCCGTGAGGAGGTGAGAGCGAAA TGAGTCCCGGCGATAGTCCGTATCCGAGATCGACGACCGTTTCGTTCCGATCCGACCCCGGCG CCGTTTTCGCACCCATTGGATGATCCCGTCGTTTACA |
| Mbox-uORF1 | GCCGGATTTGTATTAGACTAAGCTCAAGCACCTCGCTAGGTGAGGCGTCTGCGCGGATATAG GCCACTGACCTCGAACGTCGAAAGACGCCCAGGGTCAGGACAGCTCTTCCCGGCTTAAGGGT TGAGCCCAAGTGGCTTCCGGCTGGACCGGCCGGATACGCCGTGTGGTGCCAAAGCTCTGACG AGAGGGGTGCCGAGTTCGGTGGTCTGCTGGGCTGTCATCCCTTTGTGCTGTGCATCGGCATCC CCGTGTGCCCCGGCCGTGAGGAGGCATTGGATGATCCCGTCGTTTACA |

| | |
|---|---|
| Mbox-GTG-CTG$^{uORF1}$ | GCCGGATTTGTATTAGACTAAGCTCAAGCACCTCGCTAGGTGAGGCGTCTGCGCGGATATAG<br>GCCACTGACCTCGAACGTCGAAAGACGCCCAGGGTCAGGACAGCTCTTCCCGGCTTAAGGGT<br>TGAGCCCAAGTGGCTTCCGGCTGGACCGGCCGGATACGCCGTGTGGTGCCAAAGCTCTGACG<br>AGAGGGGTGCCGAGTTCGGTGGTCCATTGGATGATCCCGTCGTTTACA |
| Mbox-uORF2 | GCCGGATTTGTATTAGACTAAGCTCAAGCACCTCGCTAGGTGAGGCGTCTGCGCGGATATAG<br>GCCACTGACCTCGAACGTCGAAAGACGCCCAGGGTCAGGACAGCTCTTCCCGGCTTAAGGGT<br>TGAGCCCAAGTGGCTTCCGGCTGGACCGGCCGGATACGCCGTGTGGTGCCAAAGCTCTGACG<br>AGAGGGGTGCCGAGTTCGGTGGTCTGCTGGGCTGTCATCCCTTTGTGCTGTGCATCGGCATCC<br>CCGTGTGCCCCGGCCGTGAGGAGGTGAGAGCGAAATGAGTCCCGGCGATAGTCCGTATCCG<br>AGATCGACGACCGTTTCGTTCCGATCCGACCCCGGCGCCGTTTTCGCACTCCATTGGATGATCC<br>CGTCGTTTACA |
| Mbox-*pe20*::*lacZ* | GCCGGATTTGTATTAGACTAAGCTCAAGCACCTCGCTAGGTGAGGCGTCTGCGCGGATATAG<br>GCCACTGACCTCGAACGTCGAAAGACGCCCAGGGTCAGGACAGCTCTTCCCGGCTTAAGGGT<br>TGAGCCCAAGTGGCTTCCGGCTGGACCGGCCGGATACGCCGTGTGGTGCCAAAGCTCTGACG<br>AGAGGGGTGCCGAGTTCGGTGGTCTGCTGGGCTGTCATCCCTTTGTGCTGTGCATCGGCATCC<br>CCGTGTGCCCCGGCCGTGAGGAGGTGAGAGCGAAATGAGTCCCGGCGATAGTCCGTATCCG<br>AGATCGACGACCGTTTCGTTCCGATCCGACCCCGGCGCCGTTTTCGCACTCTGAATCGGCCTTC<br>CGGTTCGAAATCCGTTATTTCGCAAGCTCGTTGCTTCGCGGCCTTGTGTGAGTGACGTTCACG<br>GGAAGTAGCCACGACAGAAGCGGTCATAGGCCTCCGGGTTCGGTCGTCTGTCAGGAGAAGA<br>CCCATGCATTGGATGATCCCGTCGTTTACA |
